# Applying the auxin-inducible degradation (AID) system for rapid protein depletion in mammalian cells

**DOI:** 10.1101/182840

**Authors:** Bramwell G. Lambrus, Tyler C. Moyer, Andrew J. Holland

**Author notes:** These authors contributed equally to this work (Bramwell G. Lambrus, Tyler C. Moyer). Correspondence should be addressed to A.J.H. Johns Hopkins, University School of Medicine, 702A PCTB, 725 North Wolfe Street Baltimore, MD, 21205.

## Abstract

The ability to deplete a protein of interest is critical for dissecting cellular processes. Traditional methods of protein depletion are often slow-acting, which can be problematic when characterizing a cellular process that occurs within a short period of time. Furthermore, these methods are usually not reversible. Recent advances to achieve protein depletion function by inducibly trafficking proteins of interest to an endogenous E3 ubiquitin ligase complex to promote ubiquitination and subsequent degradation by the proteasome. One of these systems, the auxin-inducible degron (AID) system, has been shown to permit rapid and inducible degradation of AID-tagged target proteins in mammalian cells. The AID system can control the abundance of a diverse set of cellular proteins, including those contained within protein complexes, and is active in all phases of the cell cycle. Here we discuss considerations for the successful implementation of the AID system and describe a protocol using CRISPR/Cas9 to achieve bi-allelic insertion of an AID degron in human cells. This method can also be adapted to insert other tags, such as fluorescent proteins, at defined genomic locations.

## Introduction

The ability to deplete proteins from mammalian cells is critical for studying their role in biological processes. Traditional methods of disrupting protein function include DNA editing to produce a gene knockout (Sauer & Henderson, 1988), or RNA interference (RNAi) to downregulate mRNA (Elbashir et al., 2001). While these methods are staples of a cell biologist’s toolkit, in both cases protein depletion is indirect and the rate of protein loss depends on the stability of the protein. Furthermore, neither gene knockout nor RNAi is easily reversible, and RNAi can suffer from incomplete silencing and off-target effects (Bartlett & Davis, 2006).

To overcome the slow protein depletion achieved with genetic manipulation, researchers have turned to modulating protein function with cell-permeable, small molecules. While fast-acting, small-molecule probes are challenging to develop and frequently limited by specificity. However, combining pharmacologic manipulation with genetic engineering expands the possibilities for achieving rapid and specific modulation of protein function. One powerful example is the analog-sensitive (AS) kinase method pioneered by Shokat and colleagues (Bishop et al., 1998; K. Shah, Liu, Deirmengian, & Shokat, 1997). In this approach, mutation of conserved residues in the ATP binding pocket of a kinase allows the kinase to accommodate, and be specifically inhibited by, bulky non-hydrolyzable ATP-analogs. Importantly, wild type kinases are resistant to such inhibition. The AS kinase approach achieves specificity and reversibility, but is so far limited to manipulating proteins in the kinase family. More recent advances in chemical genetics have opened the possibility of regulating classically non-druggable targets by tagging proteins of interest with degrons (Banaszynski, Chen, Maynard-Smith, Ooi, & Wandless, 2006; Iwamoto, Bjorklund, Lundberg, Kirik, & Wandless, 2010). In these cases, proteins can be stabilized or targeted for degradation through the introduction of small molecules. A comparison of the protein degradation systems currently available is shown in Table 1.

**Table 1.** Methods for Protein Depletion in Mammalian Cells

| Methods for Protein Depletion in Mammalian Cells |  |  |  |  |
| --- | --- | --- | --- | --- |
| Name | Explanation | Advantages | Disadvantages | Ref. |
| RNAi | Small interfering RNAs (siRNAs) targeting the mRNA of a protein of interest are introduced to the cell to activate an intrinsic RNA degradation pathway. Degradation of mRNA leads to subsequent protein loss. | Only requires commercially available siRNAs. | Incomplete protein depletion<br>Slow-acting. Relies on turnover of the endogenous protein.<br>siRNAs can silence off-target mRNAs.<br>Not readily reversible. |  |
| Gene Knockout | Genomic loci are mutated through a variety of methods. Mutations lead to the introduction of premature stop codons or the generation of a non-functional protein. | Complete loss of the protein.<br>Gene-specific. | Slow-acting. Relies on turnover of the endogenous protein.<br>Not readily reversible. |  |
| Halo-Tag Degron | Protein of interest is tagged with a Halo tag, which can be bound by a hydrophobic, cell-permeable ligand. This ligand mimics an unfolded protein and targets the Halo-tagged protein for degradation. | Gene-Specific<br>Can be tuned with dosage of the synthetic ligand. | Slower degradation kinetics than the AID system.<br>Addition of the Halo tag (33 kDa) may disrupt protein function. | (Neklesa et al., 2011) |
| (DD)-FKBP12 | Protein of interest is tagged with the destabilizing domain (DD) of the FK506- and rapamycin-binding protein (FKBP12). Protein is constitutively degraded until a synthetic ligand, Shield-1, is provided that protects the protein from degradation. | Gene-specific.<br>Reversible.<br>Can be tuned with the dosage of Shield-1. | Slower degradation kinetics than the AID system.<br>Expression of normal protein levels requires the constant presence of a synthetic ligand (i.e Drug "On").<br>Addition of the DD domain (12 kDa) may disrupt protein function. | (Banaszynski et al., 2006) |
| (DD)-DHFR | Protein of interest is tagged with the intrinsically unstable modified dihydrofolate reductase (DHFR) from E.coli. Protein is constitutively degraded until the ligand, trimethoprim, is added. | Gene-specific.<br>Reversible.<br>Can be tuned with the dosage of synthetic ligand. | Slower degradation kinetics than the AID system.<br>Expression of normal protein levels requires the constant presence of trimethoprim (i.e Drug "On"). | (Iwamoto et al., 2010) |
| LID domain | Protein of interest is tagged with a Ligand-Induced Degradation (LID) domain. Addition of a small molecule, Shield-1, changes the conformation of the degron, leading to its recognition by an E3 ubiquitin ligase and subsequent degradation. | Reversible.<br>Gene-specific.<br>Can be tuned with the dosage of Shield-1. | Slower degradation kinetics than the AID system.<br>Degradation of the LID-tagged protein is often incomplete.<br>Addition of the LID domain (14 kDa) may disrupt protein function. | (Bonger, Chen, Liu, & Wandless, 2011) |
| deGradFP | Protein of interest is tagged with GFP. Expression of an F-box protein fused to a GFP-nanobody causes the protein of interest to be trafficked to the SCF E3 ligase complex for ubiquitination and subsequent degradation. | Gene-specific.<br>When introduced in a transgenic organism, the nanobody-tagged F-box protein can be placed under a tissue specific promoter. | Degradation kinetics are dependent upon expression level of the nanobody/F-box protein fusion.<br>Degradation is not inducible with a small molecule.<br>Addition of the GFP (27 kDa) may disrupt protein function. | (Caussinus, Kanca, & Affolter, 2011) |
| Auxin-Inducible Degron (AID) | Protein of interest is tagged with the auxin-inducible degron (AID). In the presence of the plant hormone auxin, the AID interacts with the F-box protein OsTIR1, bringing the protein of interest to the SCF E3 ubiquitin ligase complex for ubiquitination and subsequent degradation by the proteasome. | Rapid kinetics of protein degradation.<br>Reversible.<br>Gene-specific.<br>Can be tuned with dosage of auxin.<br>Inexpensive ligand (auxin). | Two component system requiring addition of the AID degron and expression of the F-box protein OsTIR1.<br>Addition of the mAID degron (5 kDa) may disrupt protein function. | (Nishimura et al., 2009) |
| Jasmonate-Inducible Degron (JID) | Protein of interest is tagged with the JAZ degron. In the presence of isoleucine-conjugated jasmonate (JA-Ile), the JAZ degron interacts with the F-box protein COI1, bringing the protein of interest to the SCF E3 ubiquitin ligase complex for | Fast kinetics of protein degradation.<br>Reversible.<br>Gene-specific.<br>Can be tuned with dosage of JA-Ile. | Two component system requiring addition of the JAZ degron and expression of the F-box protein (COI1).<br>Kinetics of degradation are slower than the AID system.<br>Addition of the JAZ degron (3 | (Brosh et al., 2016) |
|  | ubiquitination and subsequent degradation by the proteasome. |  | kDa) may disrupt protein function. |  |
| PROTAC | PROTAC: Proteolysis targeting chimeric molecules. Bifunctional molecules are introduced that bind to an E3 ubiquitin ligase complex in addition to a protein of interest. The protein of interest is brought to the E3 ubiquitin ligase complex, ubiquitinated, and degraded by the proteasome. | • No genetic modification required.<br>• Reversible.<br>• Gene-specific.<br>• Can be tuned with dosage of specific PROTAC. | • Applicable only to proteins that have known small molecules that bind to them.<br>• Varying kinetics depending on binding affinity of the protein and the small molecule. | (Collins, Wang, Caldwell, & Chopra, 2017; Lu et al., 2015; Winter et al., 2015) |

### Hijacking the SCF complex

Several systems have achieved protein degradation by ectopically targeting proteins to endogenous E3 ubiquitin ligase complexes, including the VHL, MDM2, cIAP, CRBN, and SCF complexes (Lu et al., 2015; Nishimura, Fukagawa, Takisawa, Kakimoto, & Kanemaki, 2009; Sakamoto et al., 2001; Schneekloth, Pucheault, Tae, & Crews, 2008; Sekine et al., 2008; Winter et al., 2015). The SCF (Skp1, Cul1, F-box) complex is a member of a family of cullin-RING-ligase E3 ubiquitin ligases (Skaar, Pagan, & Pagano, 2013). The Cul1 subunit acts as the major scaffold, bringing together Skp1 and RBX1. RBX1 is responsible for recruiting the E2 ubiquitin-conjugating enzyme to the complex, while Skp1 associates with an F-box protein that interacts with substrates containing a degradation motif (Figure 1A). The human genome encodes nearly seventy unique F-box proteins that each interact with a distinct subset of substrates, allowing the SCF complex to serve as a versatile tool for controlling the proteome (Kipreos & Pagano, 2000).

**Figure 1.**
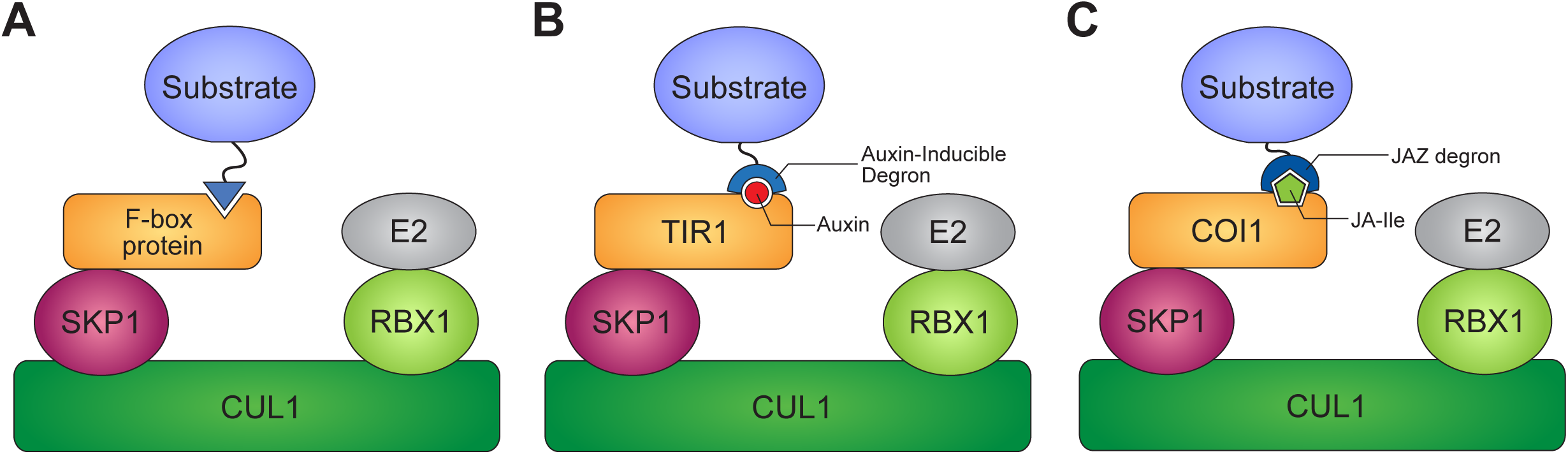
SCF E3 ubiquitin ligase complexes. A) Substrates are brought to the SCF complex through an interaction with a specific F-box protein. Due to the conserved nature of the SCF complex, F-box proteins from one kingdom can be introduced into other species to produce a functional SCF complex. B) In the AID system, the interaction between the F-box protein TIR1 and an AID-containing substrate is facilitated through the hormone Auxin. C) In the JAZ degron system, the interaction between the F-box protein COI1 and a JAZ degron-containing substrate is facilitated through the hormone jasmonate-isoleucine (Ja-Ile).

The SCF complex is highly conserved among eukaryotes (Skaar et al., 2013), making it possible to transplant F-box proteins from one organism into another, to form a functional SCF E3 ligase complex that can instruct the degradation of proteins tagged with a cognate degron. One such F-box/degron system that has been transplanted across kingdoms is the auxin-inducible degron (AID) system (Nishimura et al., 2009) (Figure 1B). In the presence of auxin (a class of structurally similar plant hormones, of which Indole-3-acetic acid (IAA) is the most common), the plant-specific F-box protein transport inhibitor response 1 (TIR1) associates with the auxin or Indole-3-Acetic Acid (AUX/IAA) family of transcription factors. This brings the AUX/IAA transcription factors to the SCF complex for their ubiquitination and subsequent degradation by the proteasome (Dharmasiri, Dharmasiri, & Estelle, 2005; Kepinski & Leyser, 2005; Tan et al., 2007).

### The Auxin Inducible Degron

Nishimura and colleagues first demonstrated the plant-specific F-box protein TIR1 could form a functional SCF^TIR1^ complex and facilitate the conditional degradation of proteins tagged with an auxin inducible degron (AID) in budding yeast and human cells (Nishimura et al., 2009). In the presence of auxin, cells expressing TIR1 can degrade an AID-tagged GFP to completion within one hour. The AID system has subsequently been used in fission yeast, *C. elegans*, and *D. melanogaster* (Kanke et al., 2011; Trost, Blattner, & Lehner, 2016; Zhang, Ward, Cheng, & Dernburg, 2015). Thus, the AID system can be used in a variety of organisms to manipulate protein function with acute temporal precision.

The AID system has additionally been shown to be active against a diverse set of cellular proteins, including those contained within macromolecular complexes, and is active in all phases of the cell cycle (Holland, Fachinetti, Han, & Cleveland, 2012). Importantly, the system is rapidly reversible, allowing AID-tagged proteins to begin re-accumulating within minutes of auxin removal. Nevertheless, a complete restoration of protein levels can take several hours and depends on the protein synthesis rate and half-life (Holland et al., 2012). In addition to the AID system, another plant-based degradation system, the JAZ degron system, has recently been transplanted into human cells (Brosh et al., 2016) (Figure 1C; Table 1). While the JAZ system has slower kinetics of protein degradation than the AID system, it nevertheless allows for the use of two orthogonal, inducible degradation systems in the same cell.

### Future optimization of the AID system

The AID system has been used to inducibly degrade a wide variety of proteins in human cells (Brosh et al., 2016; Fachinetti et al., 2015; Holland et al., 2012; Lambrus et al., 2015; McKinley et al., 2015; Natsume, Kiyomitsu, Saga, & Kanemaki, 2016; Nishimura et al., 2009). However, while many AID-tagged substrates are degraded in < 1 hour, the destruction of some abundant substrates can take longer to reach completion (Holland et al., 2012). This raises the possibility of optimizing the efficiency of AID-system to achieve more rapid and complete protein degradation. One approach is to screen for mutations in TIR1 or the AID-degron that increase the affinity of the auxin bound complex and enhance the rates of protein degradation (Yu et al., 2013). An additional possibility is to exploit the diversity of TIR1 receptors and AUX/IAA substrates across the plant kingdom. In this case, distinct TIR1 and AUX/IAA degron pairs could be adapted to tune the kinetics of protein degradation. Indeed, substrate degradation rates in yeast have been shown to vary widely depending on the specific TIR1 receptor of AUX/IAA degron motif used (Havens et al., 2012). It is worth noting that while Arabidopsis thaliana TIR1 (AtTIR1) could promote the degradation of AID-tagged proteins in yeast, it was not able to promote degradation of proteins in human cells (Nishimura et al., 2009). This is likely due to instability of AtTIR1 at 37°C, driving the need to use the thermostable TIR1 cloned from rice *Oryza sativa* to achieve efficient protein destruction in human cells (Nishimura et al., 2009).

Like other inducible degradation systems, the AID system has some capacity for “leaky” protein destruction in the absence of auxin (Natsume et al., 2016). Crystallography has shown that Auxin fills a hydrophobic cavity between TIR1 and the AID-degron to generate a stable trimeric complex interacting through a continuous hydrophobic core (Dharmasiri et al., 2005; Kepinski & Leyser, 2005; Tan et al., 2007). However, TIR1 and the AID degron can also weakly associate in the absence of auxin. In principle, one way to reduce “leaky” protein degradation would be to introduce mutations that further destabilize the TIR1-AID degron complex in the absence of auxin. An additional approach is to inducibly express the TIR1 receptor immediately prior to auxin treatment (Natsume et al., 2016).

### Tagging approaches

Several methods have been used to achieve inducible protein degradation with the AID system in mammalian cells. One approach involves the knockout or knockdown of an endogenous protein and functional replacement with an AID-tagged transgene (Brosh et al., 2016; Fachinetti et al., 2015; Holland et al., 2012). While powerful, this approach has some drawbacks, including incomplete depletion of the endogenous protein, a lack of representation of splice isoforms, and difficulty in achieving endogenous protein expression levels using a transgene. An alternative strategy that overcomes these limitations is to introduce the AID degron at the endogenous locus. This was first achieved in mammalian cells using adeno-associated viruses (AAV)-mediated gene targeting (Lambrus et al., 2015), but recently CRISPR/Cas9 genome editing has been used to more efficiently introduce the AID tag onto endogenous proteins (McKinley et al., 2015; Natsume et al., 2016).

Here, we outline a CRISPR/Cas9-based method for the generation of AID-tagged target proteins in human cell lines and describe important considerations for the successful implementation of the AID system. While we specifically focus on the use of the AID system, the scheme outlined below can also be applied to modifying endogenous loci with any tag of interest, such as fluorescent proteins, epitope tags and subcellular localization signals.

**1. Materials required**

- Expression plasmid for OsTIR1

- pBabe Neo osTIR1-9Myc (Addgene #80072)
- pBabe Blast osTIR1-9Myc (Addgene #80073)
- pBabe Puro osTIR1-9Myc (Addgene #80074)
- px459 plasmid encoding sgRNA targeting desired genomic locus

- PX459 V2.0 (Addgene #62988)
- PX459 VQR (Addgene #101715)
- PX459 VRER (Addgene #101716)
- PX459 EQR (Addgene #101732)
- PX458 (Addgene #48138)
- PX458 VQR (Addgene #101727)
- PX458 VRER (Addgene #101728)
- PX458 EQR (Addgene #101731)
- PX330 (Addgene # 42230)
- PX330 VQR (Addgene #101730)
- PX330 VRER (Addgene #101729)
- PX330 EQR (Addgene #101733)
- Plasmid template containing the mini AID tag

- pcDNA5/FRT miniAID-EGFP (Addgene #101713)
- pcDNA5/FRT EGFP-miniAID (Addgene #101714)
- Cells of choice for validation experiments and growth media for those cells
- Antibodies specific to the epitope tag of the OsTIR1 (i.e., Myc)
- Standard immunofluorescence and immunoblotting reagents
- PCR purification/concentration kit
- Transfection or nucleofection reagents

**1.1 Recipes**

- Polyethylenimine (PEI) (1 mg/mL)

- Dissolve PEI powder (25 kDa, linear) to a concentration of 1 mg/ml in water which has been heated to 80°C
- Allow solution to cool to room temp. Adjust pH to 7.0 with 5 M HCl. o Filter-sterilize the solution through a 0.22 μm membrane.
- Freeze aliquots at -80°C. Once thawed, keep at 4°C (stable for 2 months).
- Polybrene (10 mg/mL)

- Dissolve to a concentration of 10 mg/mL in water.
- Filter-sterilize the solution and freeze aliquots at -80°C. Once thawed, keep at 4°C (stable for 2 months).
- Indole-acetic acid (IAA) sodium salt (500 mM)

- Dissolve Indole-acetic acid sodium salt (Sigma # I5148) at a concentration of 500 mM in water.
- Filter-sterilize the solution and freeze aliquots in opaque tubes at -20°C. Stable at at 4°C for a few weeks. Keep protected from light.

## 2. Methods

Two modifications are required to achieve an inducible loss of protein function with the auxin-inducible degron system: 1) expression of the OsTIR1 F-box protein, and 2) integration of the AID tag into both alleles of the gene of interest. These modifications can be made in either order, though we recommend to first generate a cell line with the endogenously AID-tagged protein of interest and then to integrate the OsTIR1 transgene. High OsTIR1 expression is critical for rapid and complete degradation of AID-tagged target proteins; however, for some target proteins, very high levels of OsTIR1 expression may promote “leaky” protein degradation in the absence of auxin. Integrating the OsTIR1 transgene after AID-tagging the protein of interest offers more flexibility in identifying the optimal level of OsTIR1 expression for a specific target.

### 2.1 Designing Reagents for Site-Specific AID Integration

The CRISPR/Cas9 system is comprised of an RNA-guided nuclease that can be easily programmed to generate targeted DSBs in the mammalian genome (Wang et al., Annu. Rev. Biochem. 2016). Here, we describe how to use the well-characterized, two-component *Streptococcus pyogenes* (Sp) CRISPR/Cas9 system with engineered single guide RNA (sgRNA) for site-specific AID targeting. The specificity of SpCas9 targeting is determined by 1) Watson-Crick base pairing between the sgRNA and the genomic target sequence, and 2) direct interactions between the SpCas9 protein and a short protospacer adjacent motif (PAM) in the genomic target sequence (Figure 2A) (Mojica, Diez-Villasenor, Garcia-Martinez, & Almendros, 2009; S. A. Shah, Erdmann, Mojica, & Garrett, 2013). Reprogramming the SpCas9 to target a site is as simple as changing its guide RNA sequence. Upon recognizing a target sequence, SpCas9 cleaves the DNA 3-nt upstream of the PAM to produce a blunt-ended DSB (Figure 2A), which is then repaired either by error-prone non-homologous end-joining (NHEJ) or by homology-directed repair (HDR). For the purposes of integrating an AID tag at a specific genomic locus, we require cells to undergo HDR using a provided repair template. In the following sections, we describe how to design the guide RNA and the repair template.

**Figure 2.**
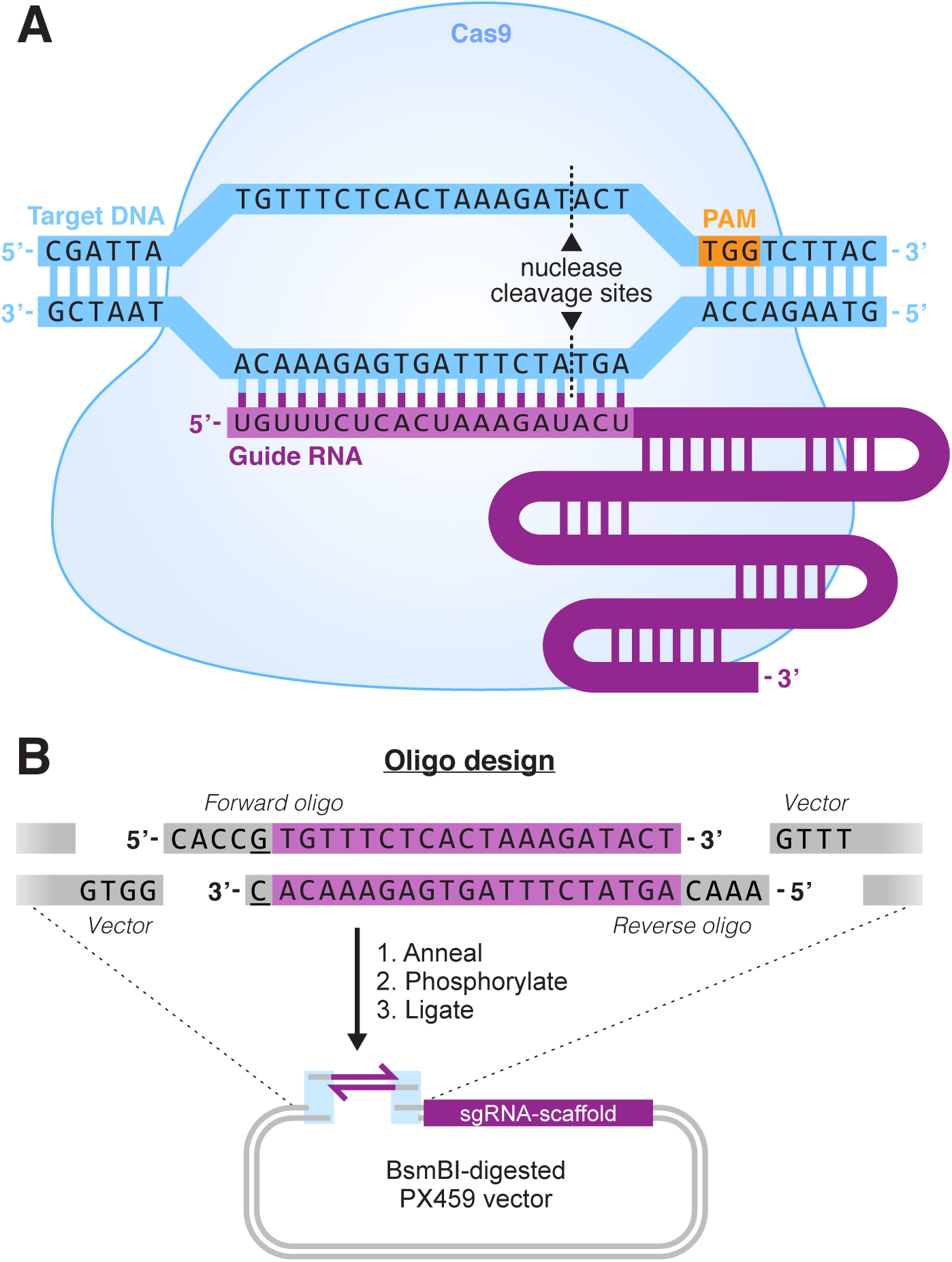
CRISPR/Cas9 system and sgRNA cloning strategy. A) Schematic of Cas9 in complex with a sgRNA, targeted to its complementary DNA sequence. Relative locations of the PAM motif (shown in orange), and nuclease cleavage sites are indicated. B) Diagram of oligonucleotide design for sgRNA cloning into PX459. A 20-nt oligonucleotide sequence is chosen, that targets the desired genomic sequence. If this sequence does not begin with a “G”, then a G should be appended to maximize expression from the U6 promoter (shown underlined). Overhangs must then be added to the oligos, with sequences complementary to those left by BsmBI digest of the PX459 vector. Oligos are then resuspended, annealed, and phosphorylated, after which they are ready for ligation into BsmBI-digested PX459.

#### 2.1.1 Choice of Cas9lsgRNA delivery system

The SpCas9 and sgRNA can either be expressed from a plasmid, or introduced into cells as a complex of purified SpCas9 protein and sgRNA (Lin, Staahl, Alla, & Doudna, 2014; Paix, Folkmann, Rasoloson, & Seydoux, 2015). Here we describe the plasmid approach, as it is the most accessible method and offers a simple means of selecting cells expressing the SpCas9 protein.

A variety of plasmids are available for SpCas9/gRNA expression in mammalian cells. Here we use the PX459 v2.0 vector (available from Addgene, #62988), which expresses SpCas9, a user-specified sgRNA and a puromycin resistance gene.

#### 2.1.2 Designing a Guide RNA for Sequence-Specific DNA Cleavage by SpCas9

The first step of sgRNA design is to determine the location of the AID tag. It is important that the tag does not impair normal localization, stability, or function of the target protein. The minimal degron, called the mini AID (mAID) is only 5 kDa, and is normally appended to the Nor C-terminus of the protein (Brosh et al., 2016). In our experience, proteins that are functional when tagged with green fluorescent protein (GFP) are also functional when tagged at the same site with mAID, though this should be empirically determined for each protein. To increase the probability of obtaining a functional AID-tagged protein, we recommend designing a strategy for tagging both the N- and C-termini of the protein.

Once the location of the AID tag has been determined, follow the steps below to design a sgRNA to direct cleavage at the specific genomic site.

1. Download the genomic sequence of the target gene from the National Center for Biotechnology Information (http://www.ncbi.nlm.nih.gov/gene).
2. Identify the position of desired AID insertion, and copy the sequence of fifty nucleotides on either side of this site.
3. Visit http://crispor.tefor.net/ (Haeussler et al., 2016) and paste the copied sequence into the query box. Next, select the appropriate genome from the drop-down menu (e.g. Homo sapiens) and select the PAM motif for the SpCas9 that is to be used (e.g. 20bp-NGG - Sp Cas9). Hit Submit.
4. The results will show a list of the potential genomic targets for sgRNA recognition, ranked by their computed “specificity score”. This score reflects the likelihood of off-target activity against other regions of the genome, and is based on the number and position of mismatches in the sgRNA (Hsu et al., 2013). High specificity scores are shown in green. While a high-scoring sgRNA is desirable to reduce the likelihood of off-target mutations, another key consideration is that HDR efficiency decreases with increasing distance from the cut site. Therefore, it is important to minimize the distance between the cut site and the desired insertion site. Taking these two parameters into account, we recommend choosing a sgRNA that has at least 4 base pair mismatches to any other sequence in the genome and promotes cutting at < 20-nt from the site where the tag is introduced. For some genomic target sequences, it is not possible to achieve these parameters, in which case we select the highest scoring sgRNA that directs cutting within 20-nt on either side of where the AID tag is to be inserted. In cases where an “NGG” PAM sequence is not available close to the site of the desired insertion, it may be possible to use PX459 expressing a modified version of SpCas9 with altered PAM specificity (SpCas9 variants available from Addgene: VQR #101715, VRER #101716, EQR #101732) (Kleinstiver et al., 2015).
5. If multiple sgRNA sequences are available for use, we recommend selecting 2-3 sgRNAs and testing each for cleavage efficiency using the SURVEYOR^®^ Mutation Detection Kit (Transgenomic), see (Ran et al., 2013).
6. To determine cloning primers for the chosen sgRNA, click “Cloning / PCR primers” under the desired guide sequence in the CRISPOR web interface. Then select the destination Addgene plasmid (pX330 and derivatives) to view the sequences necessary for cloning into PX459. Alternatively, manual oligo design is described here:

a. Copy the 20-nt sequence of the chosen genomic target sequence. The PX459 vector uses a U6 promoter to transcribe the sgRNA and this requires that a G be the first nucleotide in the transcript. In cases where the genomic target sequence does not begin with a G, append an extra G at the 5’ end of the sgRNA (Figure 2B).
b. Generate the reverse complement of the genomic target sequence (including the 5’ G if it was added) (Figure 2B).
c. Add the overhang sequence 5’-CACC-3’ to the 5’ end of genomic target and the sequence 5’-AAAC-3’ to the 5’ end of the reverse complement (Figure 2B). These sequences will produce the correct overhangs for cloning into the PX459 vector.
7. Order single-stranded DNA oligonucleotides for the two sequences generated in step 6.

#### 2.1.3 Cloning Oligonucleotides into the PX459 Vector

The PX459 vector contains two BbsI cleavage sites that allow for the insertion of annealed oligonucleotides containing the sgRNA target sequence. BbsI cleaves DNA outside of its recognition site to produce overhangs complementary to those added in step 8 above. Below we describe how to clone the sgRNA into the PX459 expression vector.

**Vector Preparation**

1. Digest 1 *μ*g of the PX459 vector with BbsI at 37°C for 2 hours.
2. To remove terminal phosphates, add 0.1 *μ*L of calf intestinal phosphatase (CIP) to the reaction and incubate at 37°C for 30 minutes.

- Since the two overhangs produced following BbsI digestion are not complementary, this step is not required, but usually reduces background.
3. Purify the cut vector using a standard PCR cleanup kit. The purified linear vector can be stored at -20°C until ready for use.

- Note that BbsI digestion of PX459 produces a small 22-nt fragment that will pass through the column, leaving a 9153-nt, linear piece of vector DNA. **Oligonucleotide Annealing**
4. Combine:

- 43 *μ*L molecular biology grade H_2_O.
- 1*μ*L of each oligonucleotide from a 100 *μ*M stock.
- 5 *μ*L of New England Biolabs Buffer 3 (www.neb.com).
5. Anneal oligonucleotides in a thermocycler with the following protocol:

a. 4 minutes at 95°C.
b. 10 minutes at 70°C.
c. Cool to 4°C at 1°C/minute. Annealed oligonucleotides can be stored at -20°C until ready for use. **Oligonucleotide Phosphorylation** We use T4 PNK from New England Biolabs to add terminal phosphates to the annealed oligonucleotides.
6. Combine:

- 5 *μ*L molecular biology grade H_2_O
- 2 *μ*L of the annealed oligonucleotides from above
- 1 *μ*L T4 PNK buffer (www.neb.com)
- 1 *μ*L ATP (10 mM stock)
- 1 *μ*L T4 PolyNucleotide Kinase (PNK) (www.neb.com)
7. Allow reaction to proceed in a thermocycler with the following protocol:

a. 30 minutes at 37°C
b. 10 minutes at 70°C (to inactivate PNK)
c. Quick cool to 4°C Phosphorylated oligonucleotides can be stored at -20°C until ready for use. **Ligation and Transformation** We use a 2X stock of Takara T4 DNA Ligase (DNA ligation kit, Version 2.1) and homemade TOP10 competent cells. In a 0.5 mL tube, combine the following:

- 1 *μ*L BbsI digested vector
- 4 *μ*L phosphorylated annealed oligonucleotides
- 5 *μ*L 2X Takara T4 Ligase
8. As a control, set up the same reaction as above but substitute 4 *μ*L molecular biology grade H_2_O for the oligonucleotides.
9. Allow ligation to proceed for one hour at room temperature.
10. Hand-thaw a frozen aliquot of competent bacteria and keep on ice.
11. Add the entire 10 *μ*L reaction mixture to the competent bacteria and incubate on ice for 20-30 minutes.
12. Heat shock the bacteria for one minute at 42°C and return to ice for at least one minute.
13. Plate the bacteria on pre-warmed ampicillin (or carbenicillin) agar plates and incubate at 37°C for 16 hours. A successful ligation should produce few (if any) colonies on a control (vector alone) plate and many fold more colonies on an experimental (vector with insert) plate.
14. Select a colony from the experimental plate and prepare a 1-5 mL culture in LB containing ampicillin. Shake at 37°C for at least 16 hours and perform a standard plasmid purification.
15. To check for correct oligonucleotide insertion, sequence the plasmid using the U6 Forward primer (5’-ACTATCATATGCTTACCGTAAC-3’).
16. PX459 plasmid DNA containing a correctly cloned sgRNA can be stored at -20°C until ready for use.

#### 2.1.4 Design of a Repair Template

To integrate AID into the site-directed DSBs generated by SpCas9, cells must undergo HDR, using a repair template containing the mAID tag flanked by homology arms to the adjacent genome sequence (Figure 3A). Repair templates can be single-stranded (ssDNA) or doublestranded DNA (dsDNA) with homology arms flanking the cut site. We recommend using a dsDNA repair template with short, 35-bp homology arms (Paix et al., 2014). In this case, the repair template can be generated by PCR-amplification of the mAID tag from a plasmid template with primers that include the homology arms (Figure 3B). We recommend include a ~10 aa flexible linker sequence between the terminus of the gene and the mAID tag to reduce the possibility of the tag disrupting protein function (Chen, Zaro, & Shen, 2013).

**Figure 3.**
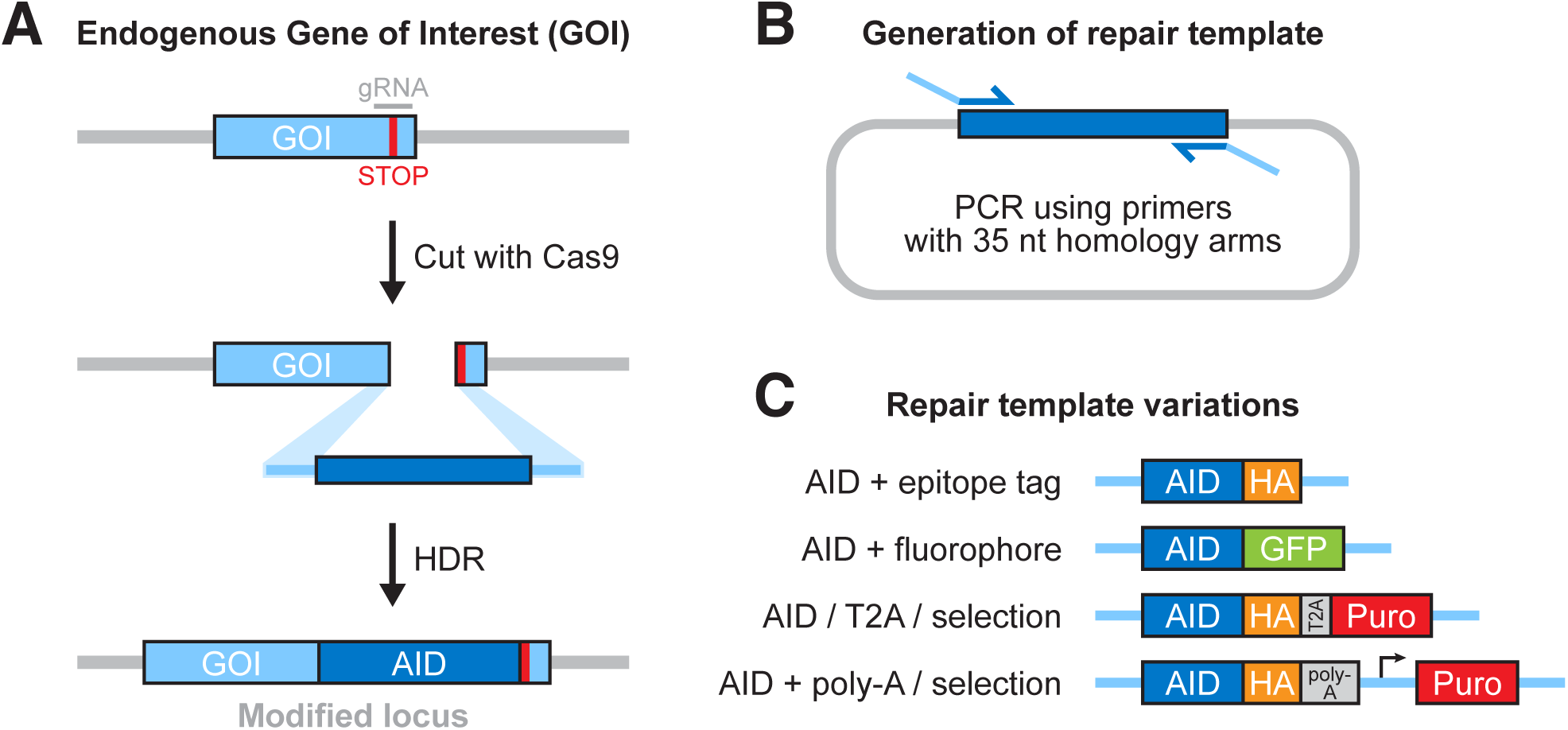
HDR and design of repair templates. A) An overview of Cas9-facilitated homology directed repair (HDR). B) A schematic showing how HDR repair templates are easily generated by PCR, using custom oligos that encode 35-nt homology arms to the gene of interest. C) Various tags and selection cassettes can be used in combination with the AID, offering different options for isolating edited cells and detecting tagged endogenous protein.

A further consideration at this point is the addition of fluorophores, tags or antibiotic resistant cassettes for downstream detection of positive clones. These tags can be appended to the AID tag by cloning the tag into the AID template plasmid prior to PCR amplification of the tag cassette. We recommend integrating an EGFP-mAID or mAID-EGFP tag, to streamline downstream identification of positive clones by fluorescence-activated cell sorting (FACS). The EGFP-mAID and mAID-EGFP template plasmid are available on Addgene (#101714 and #101713). An additional option is to include a T2A self-cleaving peptide and promoter-less antibiotic resistance gene in the repair construct, so that the tagged target protein and the selectable marker are expressed from the same transcript under the control of the endogenous gene promoter. Correctly targeted clones can then by identified by antibiotic selection. Some examples of repair constructs are shown in Figure 3C.

It is critical that the repair template should carry a mutated PAM site to prevent cutting by SpCas9 after HDR. Mutation of the “NGG” sequence is a robust way to prevent recognition by SpCas9. If it is not possible to make silent mutations that disrupt the PAM, an alternative approach is to make at least 4 silent mutations within the guide RNA-recognition sequence, preferably close to the PAM (Hsu et al., 2013). Finally, depending on the position of the cut site, tag integration itself may disrupt the PAM or sgRNA-binding site to prevent further SpCas9 recognition.

In the example below, we outline the steps for designing the PCR-amplified dsDNA repair template for introducing an N-terminal EGFP-mAID tag.

**Designing primers containing homology arms**

1. Identify the specific site of cleavage by SpCas9 in the genomic target sequence. This will be 3-nt’s upstream of the PAM sequence (Figure 2).
2. Select 35 nucleotides on either side of the cut site to act as the homology for the repair template.
3. Design primers to amplify EGFP-mAID, and append the homology arms to the primers. For example, the forward primer should consist of (5’-3’): 35-nt of the 5’ homology arm, then the EGFP-amplifying sequence (Figure 3B). **Mutation of the PAM site**
4. Identify the PAM in the repair template and replace one of the two G bases with a C or T to create a silent mutation. For SpCas9, the “NGG” PAM sequence can be mutated to anything other than “NAG” to prevent SpCas9 cleavage (Hsu et al., 2013; Jiang, Bikard, Cox, Zhang, & Marraffini, 2013). **PCR amplification of HDR template**
5. Order the finalized ssDNA primers from IDT (http://www.idtdna.com/site).
6. Follow standard PCR procedures to amplify the EGFP-mAID tag using the primers designed above, and purify the product using a PCR clean-up kit.

- Note that to achieve a highly-concentrated PCR product, we combine the products from 8 separate PCR reactions together and purify them using the MiniElute PCR purification kit from Qiagen.

### 2.2 Transfection and Screening of AID-tagged clonal lines

For genome editing experiments we recommend selecting a stably diploid cell line as this simplifies the process of achieving homozygous gene targeting. While we have successfully targeted aneuploid cell lines using the CRISPR/Cas9 system, determining the genotype of the resulting clones is more complex. Since puromycin is used to achieve rapid killing of cells that do not receive the PX459 expression vector, it is important that the chosen cell line is also puromycin sensitive. An alternative approach is to use the PX458 expression vector (available from Addgene, #48138) that co-expresses SpCas9, sgRNA and GFP, allowing fluorescent transfected cells to be directly sorted into individual wells of a 96-well plate.

There are several methods for delivering DNA to mammalian cells in culture. Here, we describe a method using Roche’s X-tremeGENE 9 transfection reagent. Depending on the cell line to be used, other DNA delivery methods (e.g. nucleofection) may be required.

**Day 1**

1. Seed cells for transfection at 2x10^5^ cells/well in 2 mL of media in a 6-well plate. Seed at least two wells per transfection, with one of these wells serving as a non-transfected control. Transfections are carried out in the presence of serum and, if desired, antibiotics. **Day 2**
2. Change media on cells 30 minutes prior to transfection.
3. In a 1.5 mL tube prepare the following for each transfected well in the order written below:

- 100 *μ*L serum-free media at room temperature
- 3 *μ*L X-tremeGENE 9 transfection reagent
- 1 *μ*g of PX459 plasmid
- ≥ 20:1 molar ratio of purified repair dsDNA template:PX459 plasmid
4. Flick tube gently ten times to mix.
5. Incubate at room temperature for 15-20 minutes.
6. Add the transfection mixture drop-wise to cells, and return cells to the incubator. **Day 4**
7. Change media on all cells (including controls) with fresh media containing puromycin (1-5 *μ*g/mL is usually sufficient). Return cells to the incubator.

- The PX459 plasmid contains a puromycin resistance marker to select for transfected cells.
8. Cells in the untransfected control wells should all die within 1-2 days in puromycin. Once control cells have died, proceed to isolate clones, described below.

- Cells should not remain in puromycin for more than 2-3 days, as the PX459 plasmid does not integrate into the genome and resistance to puromycin is lost over time.

#### Isolation of clonal lines

After puromycin selection, single cell clones should be isolated by cell sorting or limiting dilution. If inserting EGFP-mAID, sorting for green fluorescence by FACS will greatly reduce the number of colonies to screen. For non-fluorescent tags, obtain clonal lines by either cell sorting or limiting dilution. Below we outline how to obtain single clones using dilution cloning.

Cells are diluted across 96-well plates to obtain wells containing single cells. Since the clonogenic survival of cell lines varies greatly, we recommend seeding multiple 96-well plates with varying cell densities (1 cell/well, 3 cells/well, 10 cells/well and 30 cells/well). A 96-well plate that has growth in 10% of the wells will have a ~ 90% probability of a given well having a single colony.

1. Add 15 mL of puromycin-free media to a sterile reagent reservoir.
2. Add the desired number of cells to the media. For example, to achieve 10 cells/well, 1,000 cells would be added to the media.
3. Mix the cells in the media by pipetting up and down five times with a 10 mL pipette.
4. Using a multichannel pipette, add 150 *μ*L of the cell suspension to each well of the 96-well plate.
5. Repeat steps 1-4 for each cell density.
6. Wrap the 96-well plates in plastic film and return to the incubator. Allow 2-3 weeks for colonies to grow.

a. At ~10 days, colonies will be easily identifiable. We recommend to visually screen the wells for single colonies at this early stage, to be confident of clonality. At later stages, multiple colonies merge and become indistinguishable from single clones.

Allow 2-3 weeks for single cells to grow into colonies, then expand into larger wells for immunoblot analysis, or genomic DNA extraction for PCR analysis (described below). If analyzing by immunoblot, correctly targeted clones can be identified by a band shift in the endogenous protein corresponding to the size of the integrated mAID tag. Alternatively, clones that have integrated the mAID tag at the correct location can by identified by PCR analysis. In both cases, it is important to identify clones with homozygous targeting of the mAID tag. The process of extracting genomic DNA for PCR analysis is described below.

#### Genomic DNA Extraction

To extract genomic DNA from a small number of clones, we use the Sigma GenElute Mammalian Genomic DNA Miniprep Kit (G1N350). For extracting genomic DNA from larger numbers of clones we use a protocol adapted to use with 96-well plates and outlined below.

1. Using a tissue culture microscope, identify wells containing a single colony. Transfer 96 individual clones into 24-well plates. Return cells to the incubator.
2. When the clones are confluent, trypsinize cells in 200 *μ*L of 0.05% trypsin.
3. Remove 160 *μ*L of the trypsinized cell suspension and place into a single well of a 96-well plate with U-bottom wells.
4. Add 1 mL of media to the cells remaining in the 24-well plates and return plates to the incubator.
5. Spin the 96-well plate at 2000 RPM in a swinging-bucket rotor for 10 minutes to pellet cells.
6. To remove supernatant, quickly invert the plate to remove media and remove excess liquid by blotting on paper towels.
7. Resuspend cells in each well with 150 *μ*L PBS and spin at 2000 RPM for 10 minutes. Remove PBS as in step 6.
8. To lyse cells, add 50 *μ*L of lysis buffer (10mM Tris-HCL pH 7.5, 10 mM EDTA, 0.5% SDS, 10 mM NaCl, 1 mg/mL Proteinase K) and seal the plate with parafilm. Place the plate into a humidified chamber at 60°C overnight. A humidified chamber can be created by placing a few inches of water in a small plastic container with a sealable lid. The 96-well plate is placed in the sealed container on top of a test tube rack so that it rests above the water level.
9. The next day, remove the plate from the humidified chamber and cool to room temperature.
10. Add 100 *μ*L ice-cold EtOH/NaCl mix (75 mM NaCl in ~100% EtOH; forms a cloudy solution) to precipitate DNA and mix well.
11. Incubate at room temperature for 30 minutes.
12. Spin at 4000 RPM in a swinging-bucket rotor for 20 minutes to pellet precipitated DNA.
13. Decant liquid as in step 6.
14. Rinse pellet with 150 *μ*L cold 70% EtOH and spin for 10 minutes at 4000 RPM.
15. Decant liquid as in step 6.
16. Repeat washing step (#14-15) and air-dry DNA for 10 minutes.
17. Add 50 *μ*L TE pH 8.0 (10 mM Tris-HCl pH 8.0, 1 mM EDTA) to genomic DNA pellet.
18. Cover plate with parafilm and incubate at 50°C for 2 hours.

Genomic DNA is ready for further screening or may be stored at 4°C until ready for use.

#### Screening

HDR can be tracked by PCR amplification of genomic DNA. Forward and reverse genomic primers are designed to bind to regions of the genome outside of the homology arms of the repair template to amplify a region of 250 – 500-nt. Primers are designed using the Primer3 program (biotools.umassmed.edu). We recommend screening at least three different forward and reverse genomic primers in all possible combinations to identify a primer pair that yields a strong and specific PCR product from low quantities of DNA. The genomic primer pair will amplify the untagged allele, and can also amplify a modified locus that incorporated a small tag (e.g. mAID alone, or mAID-HA), which would appear as a band of increased size. If the incorporated insert is large, however, a third tag primer should also be designed that will bind to a sequence contained within the mAID tag. In combination with the opposing genomic primer, this primer will amplify a band specific for the mAID-targeted allele. The following sequence has worked for us in the past with the mAID tag: 5’-CCGCTAGACTTCTGACAGG-3’. When selecting primers, keep in mind that the tag-specific primer pair should amplify a different-sized product to that amplified from the untagged allele with the genomic primer pair (Figure 4A). In this way, homozygous untagged, heterozygous mAID-tagged, and homozygous mAID-tagged clones can be readily distinguished.

**Figure 4.**
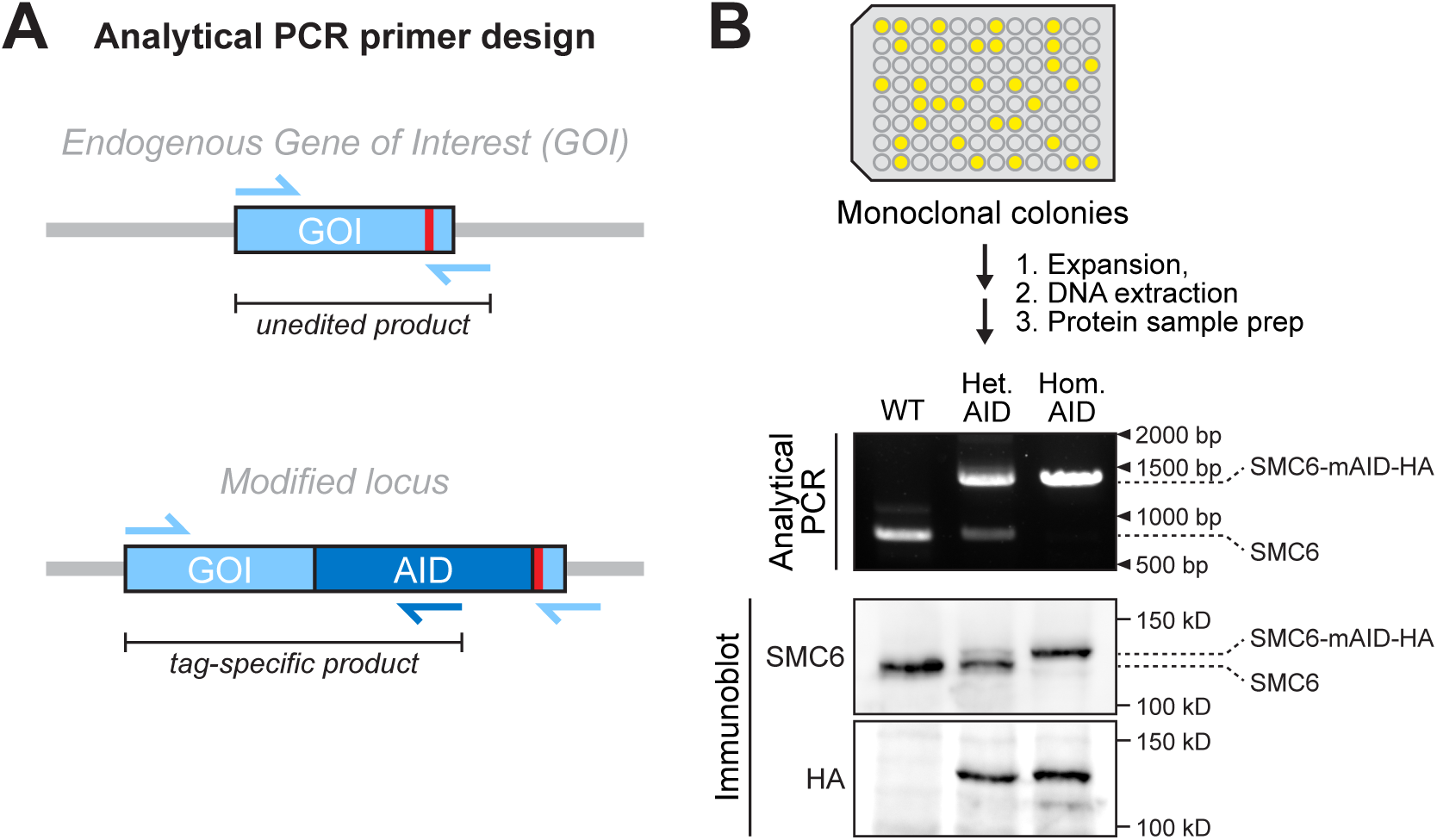
Analysis of edited cell lines. A) Analytical primers are designed to amplify a small region around the unedited, endogenous site. These primers may also be sufficient to detect modified loci that incorporate small tags (e.g. AID alone, or AID-HA), which would be observed as a product of larger size. To detect the recombination of larger inserts, however, it is recommended to design a primer within the tag itself, which would yield a tag-specific product only from edited cells. B) After expanding monoclonal colonies to enable collection of DNA and protein sample, clones can be analyzed by analytical PCR and immunoblot. The example shown here depicts results from successful editing of endogenous SMC6. PCR amplification using primers that amplify a small region around the edited site results in expected products for WT, heterozygous (Het.), and homozygous (Hom.) clones, where inclusion of the mAID-HA tag yields a larger PCR product. The SMC6 immunoblot likewise shows increased protein size from fusion of the mAID-HA tag, and the HA immunoblot shows successful tagging with HA epitope.

In most cases, the PCR protocol outlined below produces good results. However, PCR optimization may be required.

#### PCR Amplification

1. Prepare a master mix (100 reactions per plate) as follows:

- Molecular biology grade H_2_O 1060 *μ*L
- 5X GC Buffer (www.neb.com) 400 *μ*L
- Forward Genomic Primer (10 *μ*M stock) 100 *μ*L
- Reverse Genomic Primer (10 *μ*M stock) 100 *μ*L
- Forward or Reverse Tag Primer (10 *μ*M stock) 100 *μ*L
- dNTPs (stock with each dNTP at 10mM) 40 *μ*L
- Phusion Polymerase (www.neb.com) 40 *μ*L
2. Dispense 18 *μ*L of the master mix to each well of a 96-well plate.
3. Add 2 *μ*L of genomic DNA from individual clones to each well of the 96-well plate.
4. Cover the plate with a piece of sealing film and perform a PCR reaction using the following parameters:

a. 98°C 30 seconds
b. 98°C 10 seconds
c. 57 °C 15 seconds
d. 72°C 15 seconds
e. Repeat steps b-d 34X
f. 72°C 5 minutes
g. 12 °C hold
5. Run products on an agarose gel and image on a gel documentation system. Identify positive clones by the relative size of the PCR amplicons (Figure 4B).

There are three possible results for each PCR reaction.

1. Band corresponding to the size of the PCR product amplified by the untagged allele only. This reflects the size of the endogenous sequence, which did not integrate the mAID tag. It is possible that NHEJ occurred, resulting in the creation of insertions or deletions (InDels) that prevent further cutting by SpCas9.
2. Band corresponding to the size of the PCR product amplified by the tagged allele only. This clone is homozygous for insertion of the mAID tag.
3. PCR products amplified by both the tagged and untagged allele. There are two explanations for this result. 1) The clone is heterozygous. One allele integrated the tag by HDR and the other did not. In most cases the untagged allele this will have undergone NHEJ, which may generate frameshift mutations. If you are adding an amino terminal, mAID tag and the guide RNA was designed to cut in the coding sequence of the target gene, it is possible for cells to carry a tagged-allele and a frameshifted null allele. In this case, all the endogenous protein carries the mAID tag and the clone may be suitable for further analysis. 2) An alternative possibility is the cells are polyclonal. This may occur when two or more parental cells grow in a single well.

#### Sequencing Clones

To verify insertion of the desired mutation, the PCR product can be cloned into a vector for sequencing. We recommend using the ZeroBlunt^®^ TOPO^®^ Cloning Kit from Invitrogen, which allows for easy cloning of blunt end PCR products into the pCR-Blunt II-TOPO vector. We usually sequence ~10 clones to ensure sequence coverage of both alleles.

Clones that are homozygous for the mAID tag can be taken forward for further analysis. Heterozygous clones that are tagged in one allele, but carry a functional second allele, may serve as useful controls.

### 2.3 Generation of an OsTIR1 Cell Line

Once a cell line has been produced in which both endogenous alleles are tagged with a mAID degron, we then integrate the OsTIR1 protein and isolate monoclonal cell lines. An important consideration is that low expression of OsTIR1 can be rate-limiting to the degradation of AID-tagged targets, and therefore it is recommended to screen several clones by immunoblot and select a fast-growing and high-expressing OsTIR1 clone. Plasmids encoding osTIR1 are available from Addgene (Neo #80072, Blast #80073, Puro #80074).

#### 2.3.1 Production of OsTIR1 Retrovirus

1. Coat a 10 cm dish with poly-L-lysine. Incubate for 30 min at room temp. Remove and rinse once with 1x PBS.
2. Seed 3x10^6^ HEK293GP cells. Allow to settle overnight.
3. 24 hours later, prepare the following transfection cocktail:

- 600 *μ*L Opti-MEM
- 35 *μ*L (10 mg/mL) PEI
- 4.5 *μ*g pBabe OsTIR1 retrovirus plasmid
- 2-3 *μ*g VSV-G plasmid
4. Mix gently and incubate 20 minutes at RT.
5. Add dropwise to existing media on HEK293GP cells.
6. Next day, replace with fresh medium.
7. Incubate for 48 hours.
8. Collect virus-containing media and filter through a 0.45 *μ*m filter to remove cells.
9. Viral supernatant may be saved as 1 mL aliquots and snap-frozen for later use.

#### 2.3.2 Viral Transduction of OsTIR1 into Cells

1. Seed two wells of a 6-well plate with 2x10^5^ cycling cells. One of these wells will serve as a control for antibiotic selection. Allow cells to settle overnight.

- Retroviral integration requires nuclear envelope breakdown; ensure cells are cycling at the time of transduction.
2. The next day, aspirate media and add back 1 mL complete media.
3. Add 1 mL of viral supernatant to the well. To the selection control, add 1 mL complete media. Add polybrene to a final concentration of 10 *μ*g/mL, and swirl to mix. Place back in incubator and allow two days for expression.

- Some cell lines are sensitive to polybrene. To avoid toxicity, it is recommended to replace with fresh media after overnight incubation with virus.

#### 2.3.3 Selection and isolation of OsTIRl-Expressing Cells

Two days after viral transduction, cells are ready to undergo antibiotic selection.

1. Add antibiotic to transduced and control wells, and maintain selection until all control cells have died (this will take several days, depending on cell line and antibiotic used).
2. The surviving cells should express OsTIR1. If cells are sparse, provide fresh media to allow for recovery and allow several days to grow to reasonable density.

Following selection, single cell clones should be isolated and screened to identify cells expressing high levels of OsTIR1. Either single cell sorting or dilution cloning (see above) may be used to isolate single cells.

#### 2.3.5 Screening for High-Expressing OsTIRl Clones

Once clonal OsTIR1 lines have grown, they must be expanded and screened to identify a high-expressing clone. We use the pBabe OsTIR1 construct with C-terminal 9xMyc, and immunoblot against Myc to compare expression. We describe our procedure for harvesting samples, below.

1. Identify ~6-24 of the fast-growing monoclonal lines from the 96-well plates, and expand them into 12-well plates. Allow them to grow to confluence.
2. Once the 12-wells are confluent, trypsinize the cells, resuspend in 1 mL media, and transfer 700 *μ*L of the suspension to microfuge tubes.
3. Centrifuge at 500*g* for 5 min to pellet cells. Remove the supernatant and resuspend the sample in an appropriate volume of 2x Sample Buffer.
4. Boil the sample at 95°C for 10 minutes.
5. Run lysates on an SDS-PAGE gel, transfer onto nitrocellulose, then block and blot the membrane with primary antibody against Myc (Millipore Sigma; Anti-Myc Tag clone 4A6; 05-724). Once expression of OsTIR1 is visualized, select the highest expressing clone to expand for AID-tagging.

- Very high expression of OsTIR1 may interfere with the degradation of endogenous SCF substrates and slow cell cycle progression. For this reason, we isolate only the fast-growing monoclonal cell lines.

## 3. Functional Analysis

### 3.1 Validation of mAID-fusion protein in cells

As the goal of AID-tagged degradation of proteins is to test the function of the endogenous protein in its normal context, the expression, localization, and function of the mAID-tagged protein should be validated. Expression of a protein with the correct molecular weight can be validated by immunoblotting, and protein abundance can be compared to endogenous levels. Localization can be tested by standard immunofluorescence techniques. Function of the mAID-tagged protein can be evaluated by comparison to known loss-of-function phenotypes.

### 3.2 Testing inducible depletion of mAID-tagged protein

Responsiveness of mAID-fusion proteins to auxin induction can be assayed by immunofluorescence and immunoblotting. Recommended assay conditions are described below.

#### Immunofluorescence assay

1. Prepare 6x glass coverslips in a 12-well plate.
2. Seed mAID-tagged OsTIR1 cells onto coverslips. Allow cells to attach overnight.
3. Next day, prepare an auxin (IAA) timecourse: add in 500 *μ*M IAA to treat cells for a total of 5 min, 10 min, 30 min, 1 hr, and 2 hrs. Leave one coverslip as the untreated control.
4. At the end of the timecourse, rinse cells with 1x PBS, and fix for 10 min at room temperature in 4% formaldehyde (FA) in 1x PBST.
5. Proceed with standard immunostaining procedures. Probe for either endogenous protein or epitope tag, and quantify signal across conditions.

#### Immunoblotting assay

1. Seed mAID-tagged OsTIR1 cells in 6 wells of a 12-well plate, such that cells are close to confluent the next day. Allow cells to attach overnight.
2. Next day, prepare an auxin (IAA) timecourse: spike in 500 *μ*M IAA to treat cells for a total of 5 min, 10 min, 30 min, 1 hr, and 2 hrs. Leave one well as the untreated control.
3. At the end of the timecourse, add 100 *μ*L 2x SB to each well to harvest lysates for immunoblotting.
4. Boil samples at 95°C for 10 minutes.
5. Proceed to run samples following usual SDS-PAGE procedures. Blot for either endogenous protein or epitope tag, and quantify signal across conditions.

a. Protein abundance and degradation kinetics can be tuned by altering the concentration of IAA used (Lambrus et al., 2015).

## 4. Useful Tips

### 4.1 Bi-allelic tagging

- If only heterozygous clones are obtained, in which case usually the non-tagged allele has been repaired by NHEJ (and is resistant to cutting by the original sgRNA), it may be possible to design an updated sgRNA to target the non-tagged allele. Electroporating the heterozygous line with the PX459 vector encoding the new sgRNA provides another opportunity to undergo HDR integration of the mAID at the non-tagged allele.
- To enrich for bi-allelic tagging, it is possible to supply two repair templates, such as AID-EGFP and AID-mCherry. A fraction of cells will incorporate AID-EGFP at one locus and AID-mCherry at the other, and these double-positive cells can be identified and isolated by FACS.

### 4.2 IAA reagent

- We recommend using Indole-3-acetic acid sodium salt (IAA) (Sigma I5148) as it can be directly dissolved in water to a working stock of 500 mM. The non-sodium salt requires initial solubilization with ethanol and NaOH.
- IAA is light-sensitive, and stock solutions should be kept protected from light. The 500 mM aqueous stock is normally a light-tan solution and darkening of this color indicates deterioration of the stock.

## 5. Troubleshooting

### 5.1 Lack of edited clones

- Ensure transfection or electroporation protocol is effective by testing with a GFP-expressing control plasmid.
- The tagged version of the protein may not function normally and therefore cannot support viability. Perform an RNAi and add-back using the tagged version of the protein to test if it can rescue a known phenotype(s).
- Editing frequency varies by cell line; one could have to screen several hundred clones to observe editing. To improve odds, HDR can be promoted by maximizing the likelihood that cuts are generated in S/G2 phase, as NHEJ is the default repair mechanism in G1. Variations of SpCas9 plasmids are available that provide cell cycle specific expression of Cas9 or cells can be synchronized prior to electroporation (Gutschner, Haemmerle, Genovese, Draetta, & Chin, 2016; Lin et al., 2014).
- Use the SURVEYOR^®^ assay to ensure the sgRNA promotes efficient cutting of the target locus.

### 5.2 AID-tagged protein is not degraded upon induction with auxin

- Identify and test a high-expressing OsTIR1 clone. Higher expression of OsTIR1 may be required, especially if the AID-tagged protein is of high abundance.
- Test if higher concentrations of IAA will improve responsiveness.
- This is sometimes observed when a mAID-tagged transgene is expressed from a plasmid using transient transfection. Transient transfections often produce extremely high levels of protein expression that can overwhelm the SCF^OsTIR1^ complex. We recommend using stable expression of the mAID-tagged transgene.

### 5.3 AID-tagged protein is unstable

- Test if the protein is stable when tagged at the targeted terminus with a GFP tag. It is possible that tagging prevents appropriate folding or stabilizing interactions with binding partners. If so, increasing linker length may help.
- High levels of OsTIR1 expression may cause leaky degradation of AID-tagged proteins. Test a clone with lower levels of OsTIR1 or consider using a doxycycline-inducible OsTIR1 expression system.

## Conclusion

The AID system offers a rapid, inducible, and reversible system to achieve protein depletion across a range of organisms and cell types. Here we described a CRISPR/Cas9-based method for the generation of biallelic, AID-tagged genes in mammalian cells. While this method is specific to the AID degron, the repair strategy can be applied to other tags of interest, including fluorescent proteins or epitope tags. A current limitation of the approach outlined here is the relatively low frequency of HDR compared to NHEJ in most established cell lines. Future advances in the genome engineering are likely to offer methodologies for achieving higher efficiencies of HDR and biallelic tagging.

